# High-Resolution 3D Ultra-Short Echo Time MRI with Rosette k-Space Pattern for Brain Iron Content Mapping

**DOI:** 10.1101/2022.08.24.505201

**Authors:** Xin Shen, Ali Caglar Özen, Humberto Monsivais, Antonia Sunjar, Serhat Ilbey, Wei Zheng, Yansheng Du, Mark Chiew, Uzay Emir

## Abstract

**Background:** The iron concentration increases during normal brain development and is identified as a risk factor for many neurodegenerative diseases, it is vital to monitor iron content in the brain non-invasively.

**Purpose:** This study aimed to quantify *in vivo* brain iron concentration with a 3D rosette-based ultra-short echo time (UTE) magnetic resonance imaging (MRI) sequence.

**Methods:** A cylindrical phantom containing nine vials of different iron concentrations (iron (II) chloride) from 0.5 millimoles to 50 millimoles and six healthy subjects were scanned using 3D high-resolution (0.94×0.94×0.94 mm^3^) rosette UTE sequence at an echo time (TE) of 20 μs.

**Results:** Iron-related hyperintense signals (i.e., positive contrast) were detected based on the phantom scan, and were used to establish an association between iron concentration and signal intensity. The signal intensities from *in vivo* scans were then converted to iron concentrations based on the association. The deep brain structures, such as the substantia nigra, putamen, and globus pallidus, were highlighted after the conversion, which indicated potential iron accumulations.

**Conclusion:** This study suggested that T_1_-weighted signal intensity could be used for brain iron mapping.

## Introduction

Iron is a fundamental element in many normal brain physiological processes. For example, it is found in hemoglobin for oxygen transportation, involved in neurotransmitter synthesis and metabolism as a component of enzymes [1]. In addition, it is essential in mitochondrial respiration as a cofactor in iron-sulfur cluster-containing proteins [2], and myelin synthesis as a cofactor with cholesterol [3]. While iron homoeostasis in the central milieu is tightly regulated by blood-brain interfaces [4], abnormal iron accumulation in brain does happen and has been associated with many neurodegenerative diseases, including Parkinson’s [5] and Alzheimer’s [6]. In addition, age-related brain iron accumulation in healthy subjects has been reported within deep brain structures such as the substantia nigra [7], putamen [8], and globus pallidus [9] regions. Moreover, environmental and occupational exposure to toxic metals such as manganese can lead to a disrupted iron homeostasis [10, 11]. Since the iron concentration increases during normal brain development and is identified as a risk factor for many neurodegenerative diseases, it is vital to monitor iron content in the brain non-invasively.

As a non-invasive imaging modality, magnetic resonance imaging (MRI) has been widely used in measuring *in vivo* brain iron content. Because of the paramagnetic nature of the elemental iron, its presence distorts the local magnetic field. As a result, it accelerates precession, causing incoherent phase accumulation over the echo times (TE) and resulting in shorter transverse relaxation times (*T*_2_^*^). Previous *in vivo* attempts to monitor iron content in the human brain using conventional gradient echo (GRE) techniques are *T*_2_^*^ - 1/*R*_2_^*^ fitting, susceptibility-weighted imaging (SWI) [8], and quantitative susceptibility mapping (QSM) [12]. However, limited by the relatively long TEs (on the order of milliseconds (ms) or longer), the sensitivity to high iron concentrations was limited when using GRE-based techniques [13, 14]. For instance, GRE-based *T*_2_^*^ fitting resulted in low *R*_2_^*^ values if the iron concentration exceeded 25 mg/g [13]. In addition, GRE-based QSM failed to quantify iron oxide nanoparticle concentration greater than 18 millimoles (=1 mg/mL) [15].

Another feature of the paramagnetic nature of the elemental iron is the shortening of longitudinal relaxation time (T_1_) [15]. Although the short *T*_2_^*^ caused by iron accumulation leads to fast signal decay, using TEs on the order of microseconds (e.g., 20 μs), the majority of the signals can be retained. With an appropriate flip angle and repetition time (TR), the signal intensity after reconstruction can be shifted to a T_1_-weighted image, which results in higher signal intensity originating from iron-rich phantoms and/or *in vivo* tissues than regions without iron accumulation [16]. The iron-related hyperintense signals (i.e., positive contrast) can be detected by ultra-short echo time (UTE) and zero echo time (ZTE) with sweep imaging with Fourier transformation (SWIFT) sequences.

UTE and ZTE sequences have been widely applied in detecting and tracking high iron concentrations, including preclinical and *in vivo* scans. Examples include: 1) tracking iron-labeled mesenchymal stem cells after injection into mice with a SWIFT sequence [16] and after injection into sheep with a UTE sequence [17]; 2) detection of receptor-targeted iron oxide nanoparticles in a mouse tumor model with a UTE sequence [18]; 3) UTE-based *R*_2_^*^ fitting to accurately quantify *in vivo* high liver iron concentration [13, 14]; 4) UTE-based QSM to detect phantom containing high iron concentration [19]; 5) simultaneously measuring demyelination in white matter (WM) and iron deposition in gray matter (GM) caused by multiple sclerosis with inversion recovery UTE [20]. However, most of the UTE and ZTE sequences for iron content detection sampled data based on a radial k-space trajectory, which may not be optimal in terms of peripheral k-space coverage and incoherence [21].

Noticeably, some studies have applied k-space trajectories with more curvature for iron content detection, e.g., 3D cones, which showed potential for acceleration [17, 19]. Rosette k-space trajectories sample data with high curvature per spoke, which may provide more efficient sampling in iron concentration measurement. In addition, the rosette k-space pattern is more incoherent than radial patterns [22]. Therefore, the reconstructed images with undersampled rosette trajectories may maintain image quality without suffering severe artifacts and/or noise amplification, providing a fast high-resolution acquisition strategy. However, 3D rosette k-space patterns have not yet been demonstrated in the UTE application of brain iron content detection.

This study aimed to measure brain iron concentration in healthy subjects by 3D high-resolution UTE (TE=20μs, 0.94 mm isotropic voxel) sequence with an undersampled rosette k-space pattern for fast acquisition. Phantoms prepared with different amounts of elemental iron were scanned to investigate the relationship between signal intensity by UTE sequences and iron concentration. In addition, the signal intensity within the deep brain structures such as the substantia nigra, putamen, and globus pallidus was compared to other brain tissues to discover the effect of the accumulated iron in UTE images.

## Material and Methods

The 3D rosette k-space pattern is shown in Figure 1, and described in the following equations [23]:

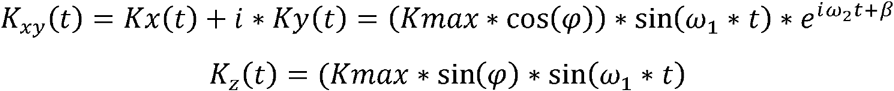

where *Kmax* is the maximum extent of k-space, *ω*_l_ is the frequency of oscillation in the radial direction, *ω*_2_ is the frequency of rotation in the angular direction, *φ* determines the location in the z-axis, and *β* determines the initial phase in the angular direction.

**Figure 1.**
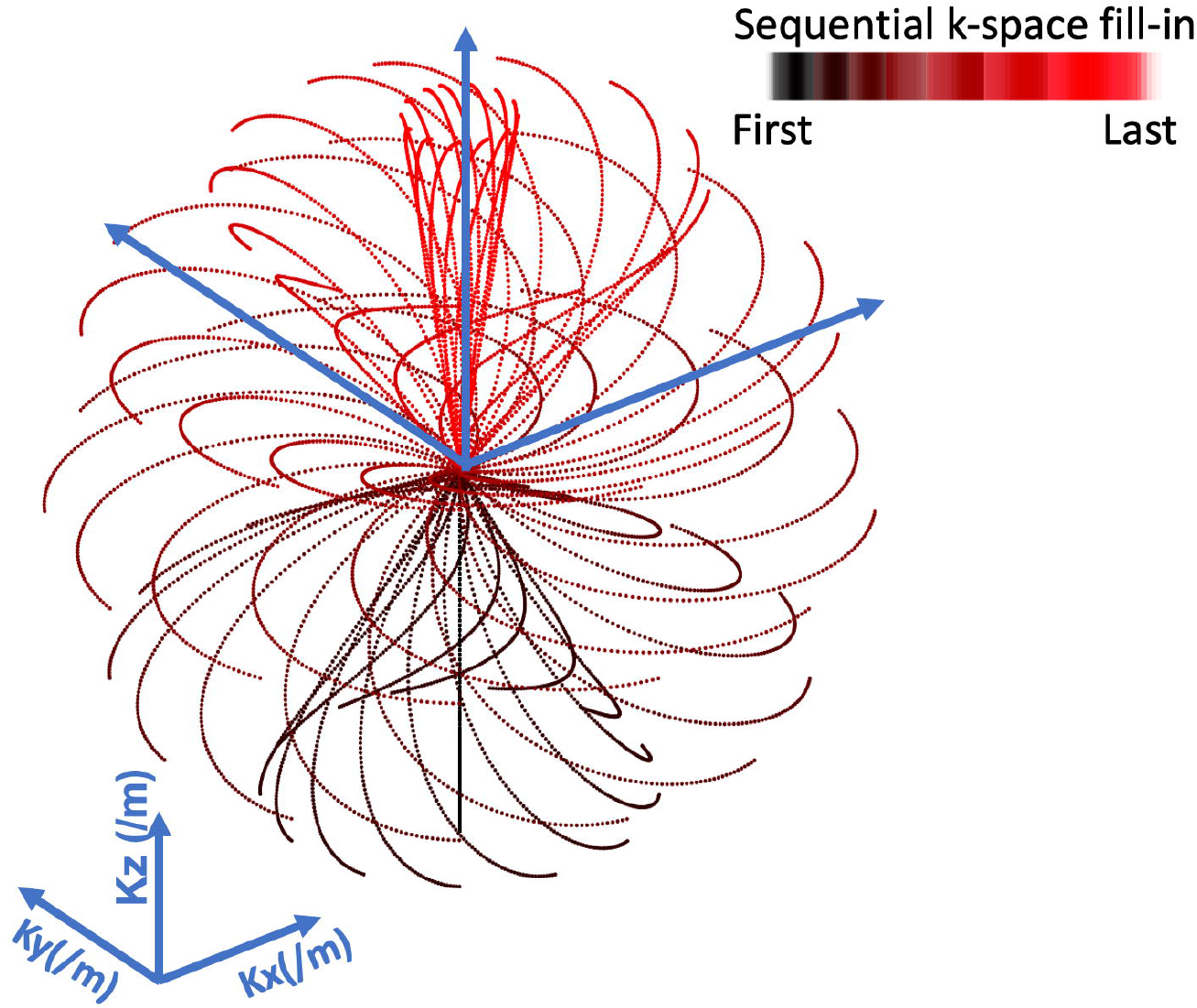
Illustration of the 3D rosette k-space design. Selective spokes with varied rotations in the k_x_-k_y_ plane and varied extensions in the k_z_-axis are shown. Each spoke samples starting from k-space origin to the maximum extension (Kmax). The 3D k-space was filled from the bottom (negative k_z_) to the top (positive k_z_).

The study was approved by the Institutional Review Boards (IRBs) of Purdue University and informed consent was obtained. All scans were performed with a whole-body 3T MRI system (Siemens Healthineers, Erlangen, Germany). A vendor-supplied 20-channel receiver head coil was used for all scans. A cylindrical phantom containing nine 15 mL vials of different concentrations of iron chloride (FeCl_2_) was constructed. A stock solution of FeCl_2_ dissolved in distilled water was diluted with an agarose solution (1% by weight) to make vials with iron concentrations of 0.5, 1.0, 2.0, 3.5, 5.0, 7.5, 10.0, 25.0, and 50.0 millimoles (mM). After the vials were fixed in a custom-made acrylic insert and submerged in the same agarose solution, the phantom was scanned using the 3D rosette UTE sequence with TE=20 μs [23]. In addition, six healthy volunteers were recruited for 3D high-resolution rosette UTE scans at TE=20 μs. The parameters for the 3D rosette UTE acquisition were: *Kmax*=500/m, *ω*_l_=*ω*_2_=0.766 kHz, number of total petals= 71442 (40% undersampling compared to 1.8 × 10^5^ required by the Nyquist criteria), samples per petal=213, *φ* was sampled uniformly in the range of [−*π*/2, *π*/2], and *β* was sampled uniformly in the range of [0,2*π*], the field of view (FOV)=240×240×240 mm^3^, matrix size=256×256×256, readout dwell time=10 *µs*, flip angle=7-degree, TR=7 ms, RF pulse duration=10 μs, readout duration=2.1 ms per echo. The total scan time was 8.7 minutes.

Image reconstruction and post-processing steps were performed in MATLAB (MathWorks, USA). The non-uniform fast Fourier transform (NUFFT) [24] and a sparsity constraint on total image variation [25] were used for image reconstruction. After image reconstruction, all images from phantom and *in vivo* scans went through 3dunifize (AFNI) [26] for bias-field correction. In addition, the *in vivo* images underwent brain tissue extraction [27] and then were registered to a standard brain atlas (MNI-152). For the phantom images, the signal intensity of different iron concentrations was quantified based on mean values of cylinder volumes of interest (VOIs) located at the centers of vials. An exponential model fitting established the relationship between iron concentration and signal intensity. The fitting model was expressed as the equation:

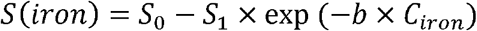

where *c*_*iron*_ represents the iron concentration in the unit of mM.

Each individual *in vivo* image was corrected using the mean signal intensity of total white matter (WM) as a marker. Assuming the mean signal intensities of the total WM of six individuals are *WM*_*n*_, where *n* represents the individual number and *n* = 1,2,3,4,5,6, the correction factor (CF) was calculated as:

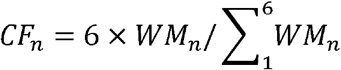

The signal intensities of all voxels were then multiplied by the corresponding individual CF. The correction aimed to minimize individual differences’ influence on signal intensity by having a constant mean signal intensity of total WM across subjects. After the correction, the *in vivo* brain iron content was converted from signal intensity directly based on the exponential relationship established in the phantom measurements. In addition, the signal intensity in several selective ROIs, including left and right substantia nigra, left and right putamen, left and right globus pallidus, cerebellum nuclei, total WM, and total GM, was measured. One-way analysis of variance (ANOVA) was performed to test the significance of signal intensity difference among ROIs. Tukey’s honest significance test (HSD) was used for multiple comparisons to find all possible pairs of means that are significantly different.

## Results

Figure 2A shows a slice of the reconstructed image from phantom scan with nine vials of different iron concentrations. Figure 2B indicates the relationship between iron concentration and VOI-based signal intensity. The signal intensity increases with higher iron concentration, following an exponential shape. The fitting result is:

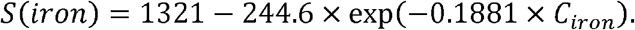

**Figure 2.**
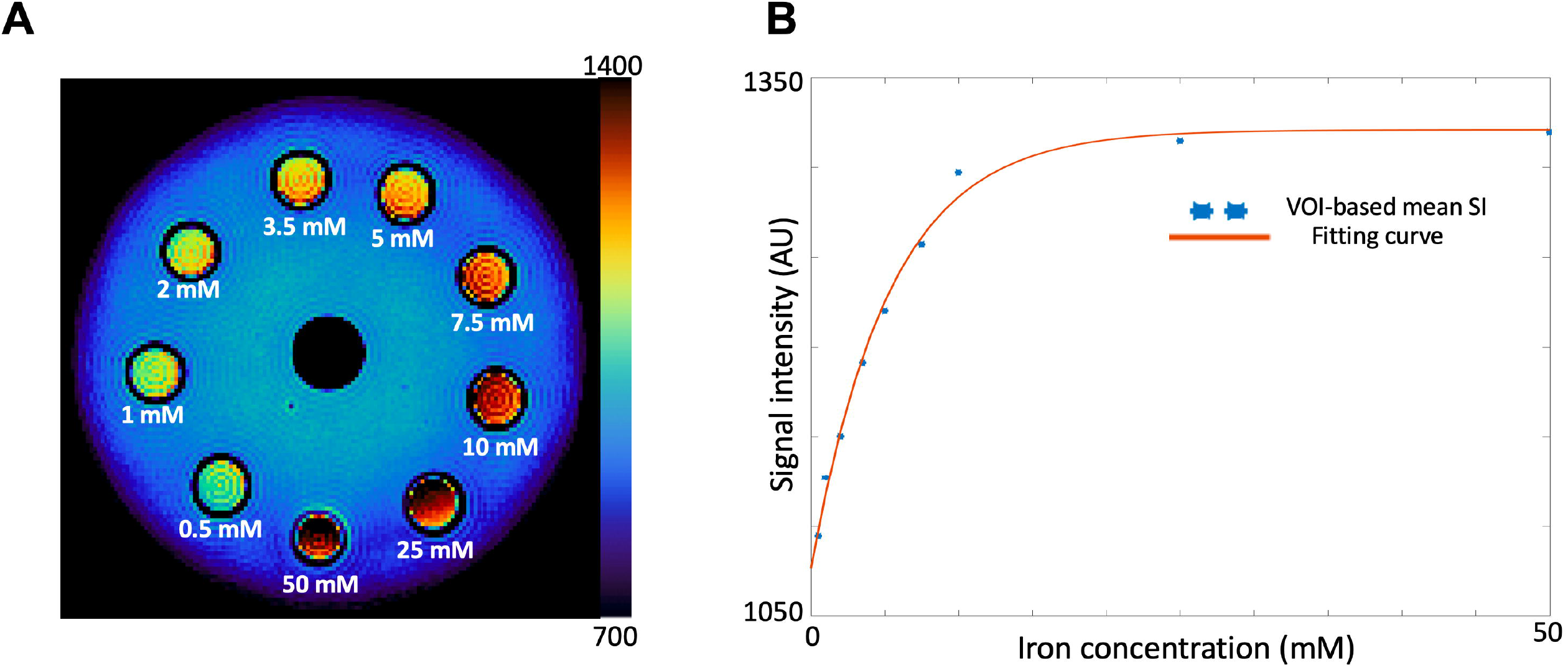
The relationship between iron concentration and the signal intensity from rosette UTE acquisition. A) The reconstructed image of a cylinder phantom containing 9 vials with different iron concentration. B) The fitting curve between signal intensity and iron concentration based on volume of interest (VOI).

The signal intensity is saturated at approximately 25 mM of iron.

Table 1 summarizes the signal intensity of six individual subjects, quantified based on multiple ROIs, including left and right globus pallidus (GP (L) and (GP (R)), left and right substantia nigra (SN (L) and SN (R)), left and right putamen, cerebellum nuclei (CN), total GM, and total WM. All results are reported as mean ± standard deviation. The *p*-values from statistical analysis reject that the signal intensity of those ROIs is from populations with the same mean value for all subjects. The highest signal intensity is reported in either left or right substantia nigra, while the lowest is in total WM.

**Table 1.** The signal intensity (mean±standard deviation) of six individuals in different ROIs, including left and right substantia nigra (SN), left and right putamen, left and right globus pallidus (GP), cerebellum nuclei (CN), total WM, and total GM.

|  | Subject 1 | Subject 2 | Subject 3 | Subject 4 | Subject 5 | Subject 6 |
| --- | --- | --- | --- | --- | --- | --- |
| GP (L) | 976 ± 49 | 981 ± 38 | 975 ± 23 | 972 ± 21 | 968 ± 36 | 967 ± 36 |
| GP (R) | 984 ± 33 | 991 ± 32 | 978 ± 22 | 972 ± 20 | 973 ± 24 | 982 ± 20 |
| SN (L) | 1070 ± 39 | 997 ± 44 | 995 ± 51 | 981 ± 55 | 1007 ± 53 | 1009 ± 51 |
| SN (R) | 1043 ± 50 | 967 ± 48 | 1008 ± 39 | 967 ± 51 | 997 ± 44 | 974 ± 37 |
| Putamen (L) | 944 ± 73 | 933 ± 96 | 933 ± 79 | 944 ± 65 | 943 ± 62 | 945 ± 69 |
| Putamen (R) | 970 ± 34 | 957 ± 54 | 953 ± 28 | 959 ± 48 | 966 ± 29 | 964 ± 46 |
| CN | 969 ± 31 | 955 ± 28 | 947 ± 28 | 958 ± 23 | 959 ± 26 | 849 ± 30 |
| GM | 964 ± 57 | 953 ± 56 | 949 ± 53 | 953 ± 70 | 957 ± 51 | 951 ± 52 |
| WM | 938 ± 43 | 938 ± 37 | 938 ± 35 | 938 ± 44 | 938 ± 40 | 938 ± 36 |
| *p*-value | < 0.0001 | < 0.0001 | < 0.0001 | < 0.0001 | < 0.0001 | < 0.0001 |

Table 2 indicates the *p*-values from Tukey’s HSD for multiple comparisons, with the mean signal intensity of different ROIs across subjects. All pairs of means are significantly different from each other, which report low *p*-values (<0.0001), except the total GM vs. cerebellum nuclei (CN) (*p*-value=0.04).

**Table 2.** The p-values from Tukey’s HSD for multiple comparisons.

|  | GP (L) | GP (R) | SN(L) | SN (R) | Putamen (L) | Putamen (R) | CN | GM | WM |
| --- | --- | --- | --- | --- | --- | --- | --- | --- | --- |
| GP (L) | 1 |  |  |  |  |  |  |  |  |
| GP (R) | < 0.0001 | 1 |  |  |  |  |  |  |  |
| SN (L) | < 0.0001 | < 0.0001 | 1 |  |  |  |  |  |  |
| SN (R) | < 0.0001 | < 0.0001 | < 0.0001 | 1 |  |  |  |  |  |
| Putamen (L) | < 0.0001 | < 0.0001 | < 0.0001 | < 0.0001 | 1 |  |  |  |  |
| Putamen (R) | < 0.0001 | < 0.0001 | < 0.0001 | < 0.0001 | < 0.0001 | 1 |  |  |  |
| CN | < 0.0001 | < 0.0001 | < 0.0001 | < 0.0001 | < 0.0001 | < 0.0001 | 1 |  |  |
| GM | < 0.0001 | < 0.0001 | < 0.0001 | < 0.0001 | < 0.0001 | < 0.0001 | 0.0040 | 1 |  |
| WM | < 0.0001 | < 0.0001 | < 0.0001 | < 0.0001 | < 0.0001 | < 0.0001 | < 0.0001 | < 0.0001 | 1 |

Figure 3 shows the transverse slices of six individual subjects after being registered to a standard brain atlas (MNI-152), containing the regions of putamen and globus pallidus. Based on the reconstructed images, there is little contrast between the putamen or globus pallidus and other brain regions. However, the difference is amplified after converting from signal intensity to iron concentration based on the established exponential relationship. Figure 4 shows the coronal slices of six individual subjects in the MNI-152 atlas, highlighting the region of the substantia nigra. Figure 5 shows the sagittal slices containing the cerebellum’s deep nuclei. Similar to the putamen, globus pallidus, and substantia nigra, the contrast between cerebellum nuclei and other brain regions was increased after the conversion.

**Figure 3.**
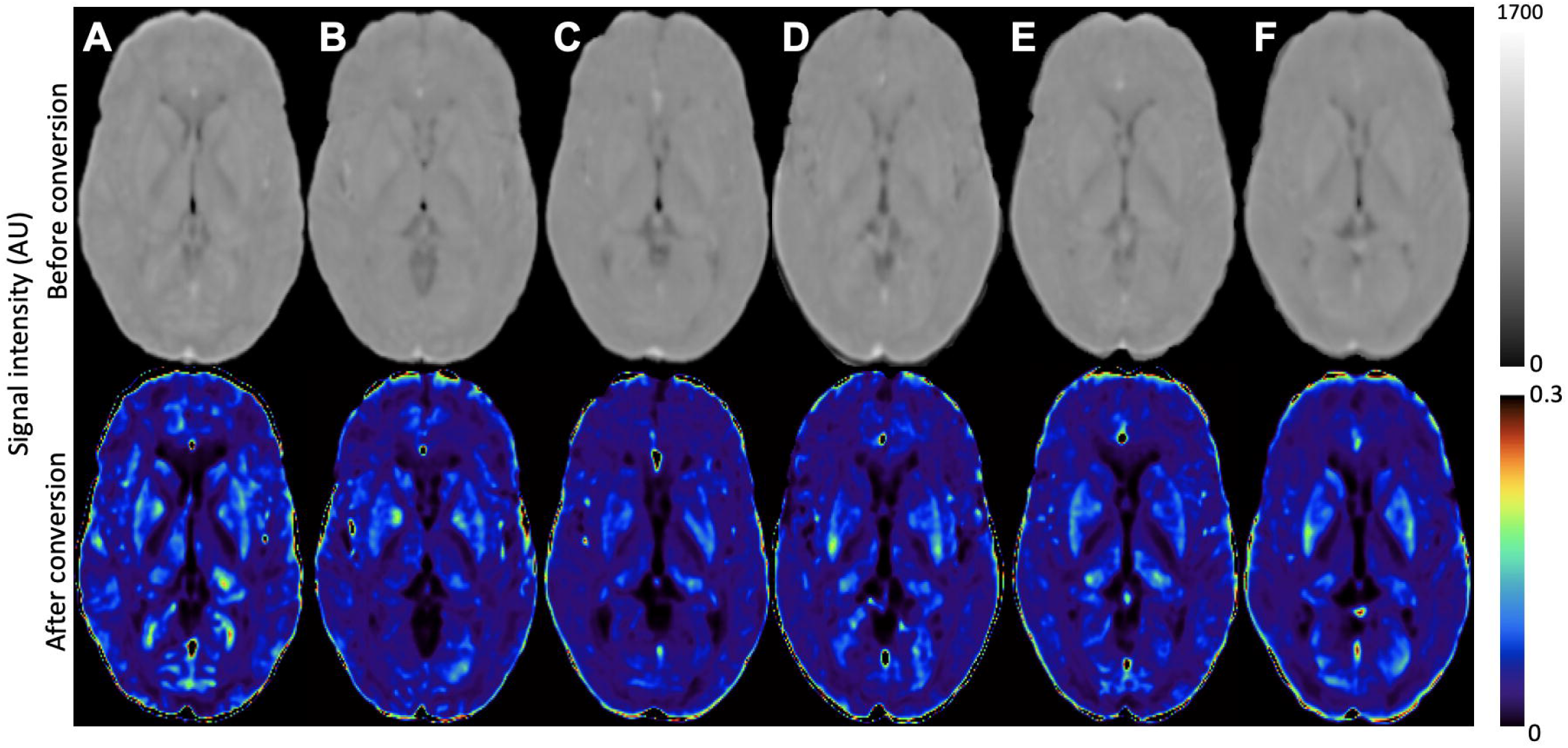
The transverse slices of six individual subjects containing the regions of putamen and globus pallidus. The top row showed the original reconstructed images, and the bottom row showed the value after conversion to iron concentration.

**Figure 4.**
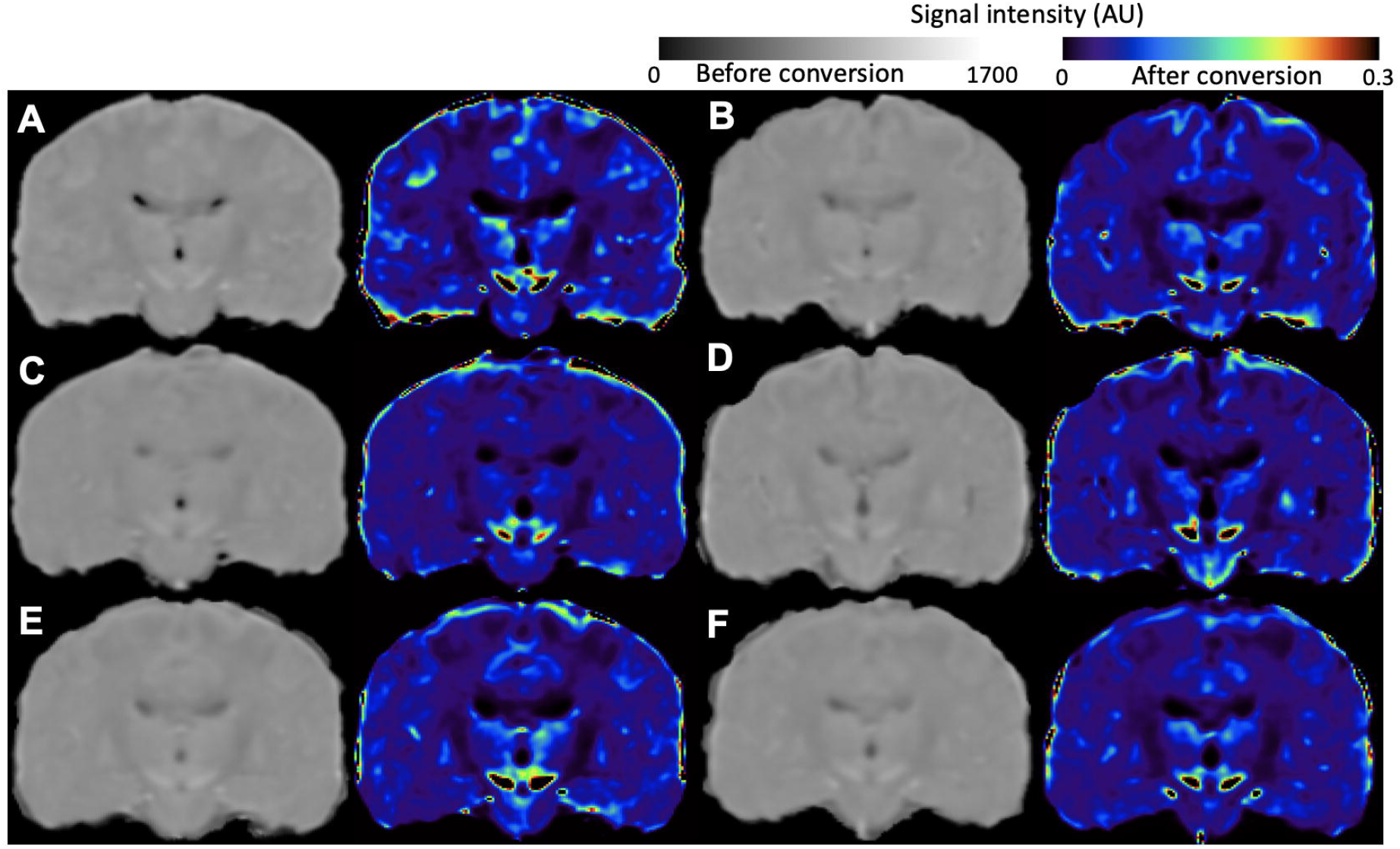
The coronal slices of six individual subjects containing the region of substantia nigra. A, B, C, D, E, F) Six individuals. Left images are from reconstruction, right images are after conversion to iron concentration.

**Figure 5.**
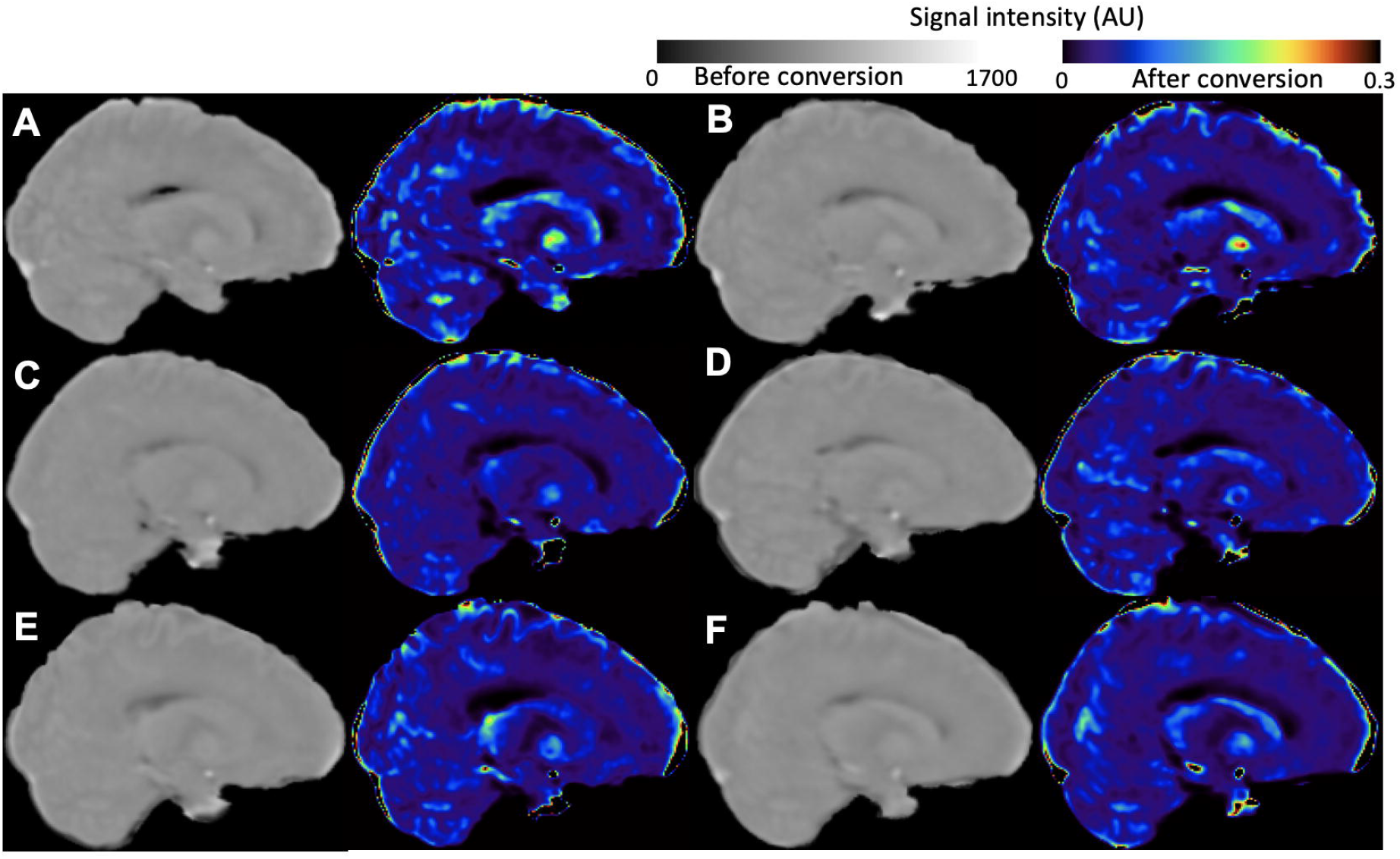
The sagittal slices of six individual subjects containing the cerebellum’s deep nuclei. A, B, C, D, E, F) Six individuals. Left images are from reconstruction, right images are after conversion to iron concentration.

## Discussion and Conclusion

This study applied a novel 3D UTE sequence based on a rosette k-space trajectory for iron content mapping. A high spatial resolution image (0.94×0.94×0.94 mm^3^) was achieved within 8.7 minutes by performing 40% undersampling strategy (relative to Nyquist criteria). A phantom scan indicated a positive contrast associated with iron concentration. The signal intensity from *in vivo* scans was converted to iron concentration with the established association. Higher iron concentration than other regions (i.e., GM and WM) was found in brain regions of the putamen, globus pallidus, substantia nigra, and deep nuclei in the cerebellum.

The iron-related MR signal intensity changes can be separated into T_1_ and T_2_* weighting effects. Recent publications on brain iron mapping used T_2_* values or T_2_*-weighted signal intensities to represent iron concentrations [13, 14]. With high iron accumulation, the T_2_* becomes too short to be detected by the conventional GRE sequence. Applying the UTE technique would overcome this problem by capturing the rapid decaying signals. However, a pure T_2_* weighting signal intensity is difficult to achieve with an ultra-short TE. This is because the T_1_ weighting of the signal intensity is hard to eliminate, especially when the accumulated iron reduces the T_1_ value. This study used T_1_-weighted signal intensity instead of T_2_*-weighted to establish an association with iron concentration. An ultra-short TE (20 *µ*s) was applied to minimize the T_2_* effect of the signal intensity, and a positive contrast was discovered.

A possible explanation for the positive contrast pertains to the decreased T_1_ value caused by the accumulation of elemental iron. A steady state is reached after several RF pulses with a short TR< T_1_ (TR=7 ms in this study). The steady-state longitudinal magnetization is highly related to TR, T_1_, and flip angle values. Therefore, a shorter T_1_ value would increase steady-state longitudinal magnetization, contributing to the positive contrast. However, an appropriate flip angle and a short TE are needed to enlarge the positive T_1_-weighted contrast.

Another factor that influences the signal intensity is the proton density (PD), which is not ruled out in this study. The PD in GM is reported to be slightly higher than in WM. However, the signal intensities within putamen, globus pallidus, substantia nigra, and deep nuclei in the cerebellum are higher than WM and GM. This indicates that PD may not be the main factor causing the high signal intensity.

This study used the mean signal intensity of total WM as a marker to generate a correction factor. This is because WM is supposed to contain less elemental iron than GM [28]. Previous studies reported an increased WM iron deposition in patients with WM injury, such as multiple sclerosis, using the magnetic field correlation technique [28] [29]. However, those inhomogeneity changes in some WM regions, such as genu of corpus callosum and frontal white matter, are small and not statistically significant [29]. On the other hand, the disease-related iron deposition changes were more noticeable and more significant in GM regions, such as globus pallidus, putamen, and thalamus, than in other brain regions [28, 29].

A potential future direction is to use T_1_ values directly to represent the iron concentration. This can be achieved by repeating scans with different flip angles. However, the TE should be ultra-short to rule out the T_2_* weighting, limiting the RF pulse duration. A selection with a wide range of flip angles may not be feasible with the current 10 *µ*s RF pulse duration. The major limitations of this study are that no comparison was performed to the brain iron quantification methods depending on T_2_* weighting and no patients with abnormal iron accumulation were recruited. Future work plans to recruit welders, who are exposed to welding fumes containing iron, for multiple MRI-based modalities for brain iron mapping.

In summary, this study showed the promising feasibility of using T_1_-weighted signal intensity for brain iron mapping. With the positive exponential relationship, the signal intensities were converted to iron concentrations. The deep brain structures, which have potential iron accumulation, were highlighted after the conversion.

## Author Contributions

Uzay Emir, Xin Shen, Wei Zheng, Yansheng Du, Mark Chiew, and Ali Caglar Ozen conceived the idea; Uzay Emir, Ali Caglar Ozen, Serhat Ilbey, and Xin Shen designed the experiments; Humberto Monsivais, Antonia Sunjar, Serhat Ilbey, and Xin Shen performed the experiments; Humberto Monsivais, Antonia Sunjar, Serhat Ilbey, and Xin Shen analyzed the data; Xin Shen, Ali Caglar Özen, Humberto Monsivais, Antonia Sunjar, Serhat Ilbey, Wei Zheng, Yansheng Du, Mark Chiew, and Uzay Emir wrote the article.

## Acknowledgments

Not applicable

